# An artificial intelligence system reveals liquiritin inhibits SARS-CoV-2 by mimicking type I interferon

**DOI:** 10.1101/2020.05.02.074021

**Authors:** Jie Zhu, Yong-Qiang Deng, Xin Wang, Xiao-Feng Li, Na-Na Zhang, Zurui Liu, Bowen Zhang, Cheng-Feng Qin, Zhengwei Xie

## Abstract

The pandemic COVID-19 has spread to all over the world and greatly threatens safety and health of people. COVID-19 is highly infectious and with high mortality rate. As no effective antiviral treatment is currently available, new drugs are urgently needed. We employed transcriptional analysis to uncover potential antiviral drugs from natural products or FDA approved drugs. We found liquiritin significantly inhibit replication of SARS-CoV-2 in Vero E6 cells with EC_50_ = 2.39 μM. Mechanistically, we found liquiritin exerts anti-viral function by mimicking type I interferon. Upregulated genes induced by liquiritin are enriched in GO categories including type I interferon signaling pathway, negative regulation of viral genome replication and etc. In toxicity experiment, no death was observed when treated at dose of 300 mg/kg for a week in ICR mice. All the organ indexes but liver and serum biochemical indexes were normal after treatment. Liquiritin is abundant in licorice tablet (~0.2% by mass), a traditional Chinese medicine. Together, we recommend liquiritin as a competitive candidate for treating COVID-19. We also expect liquiritin to have a broad and potent antiviral function to other viral pathogens, like HBV, HIV and etc.

## Introduction

Began at the end of year 2019, a novel coronavirus named SARS-CoV-2 caused outbreaks of pulmonary diseases (COVID-19) worldwide. It has greatly threatened the public health in global and killed tens-of-thousands of people^1^. By May 2nd, there are > 340,000 cumulative cases globally, with > 240,000 deaths. At present, some drugs are considered modestly effective in treating COVID-19, including chloroquine and remdesivir. However, the efficacy and safety of these drugs for SARS-CoV-2 pneumonia patients are not conclusive after several clinical trials. Current studies showed controversial effects in different trials^2^. As there is no specific treatment against COVID-19, identifying effective and safe antiviral agents to combat the disease is urgently needed.

Previous strategies for anti-viral drug development mainly focus on blocking receptors or inhibiting proteases. However, boosting the protective ability of the host was rarely considered. Herein, we consider a method to inhibit the gene expression of multiple factors that related to viral replication with one single compound. Previously, the chemical treated cell cultures were measured and analyzed by microarray or L1000 techniques^3,4^. The gene set enrichment analysis (GSEA) or Venn diagram were applied to connect the gene expression profile to up / down gene sets of certain diseases. Such technique, named connectivity map (CMAP) has been applied in screening drugs for obesity^5,6^. We proposed that similar strategy can be applied to inhibit host machinery proteins that supporting the viral replication and to induce genes that blocking the viral replication.

Liquiritin is one of the main flavonoids in *Glycyrrhiza uralensis*, which acts as an antioxidant and has antidepressant, neuroprotective, anti-inflammatory and therapeutic effects on heart system diseases^7–11^. Liquiritin plays a strong protective effect on vascular endothelial cells in myocardial ischemia-reperfusion injury model^11^. It can also protect smoking-induced lung epithelial cell injury^12^.

Type I interferon (INF) is important cytokines to protect host from viral infection. Almost all human cells can produce INFα/β, which are the best-defined type I INFs. INFs are able to induce the downstream IFN stimulated genes (ISGs) to inhibit the viral replication, including MX1, PKR, OAS, IFITM, APOBEC1, TRIM and etc^13^.

In present study, we first predicted the efficacy of compounds to inhibit the viral replication. Then, we tested them using Vero E6 cell lines. We found liquiritin is able to inhibit infection of SARS-CoV-2 efficiently with EC_50_ = 2.39 μM. Mechanistically, we found liquiritin mimicked type I INF to induce ISGs and thus protect cells from infection. We also evaluated the bioavailability, metabolism and safety of liquiritin.

## Results

### Prediction of efficacy against host genes related to viral replication

To perform GSEA against chemical induced transcriptomes, a gene set is required. In the case of SARS-CoV-2, a reasonable gene set that are enriched in viral processes by analyzing the ACE2-expressed AT2 cells were reported by Zhao et. al^14^. These genes include TRIM27, IFITM3, TMPRSS2, LAMP1 and etc. The full list is shown in Methods section. We chose the FDA approved drugs and natural products as the compound library (3682 in total). We employed InfinityPhenotype^15^, which is an artificial intelligence based platform, to predict efficacy of potential drugs (Figure 1a). InfinityPhenotype takes molecule formula as input and calculate the enrichment score, similar as in CMAP, but only one side is considered. All the scores were plotted in Figure 1b. The top scored compounds are shown in Supp. Table 1. In the natural product list, liquiritin (−0.32) and agnuside (−0.31) have emerged. In FDA approved drugs, we obtained procaterol (−0.23), pibrentasvir (−0.23) and carbocisteine (−0.21). Liquiritin ranks first among all the compounds.

**Figure 1.**
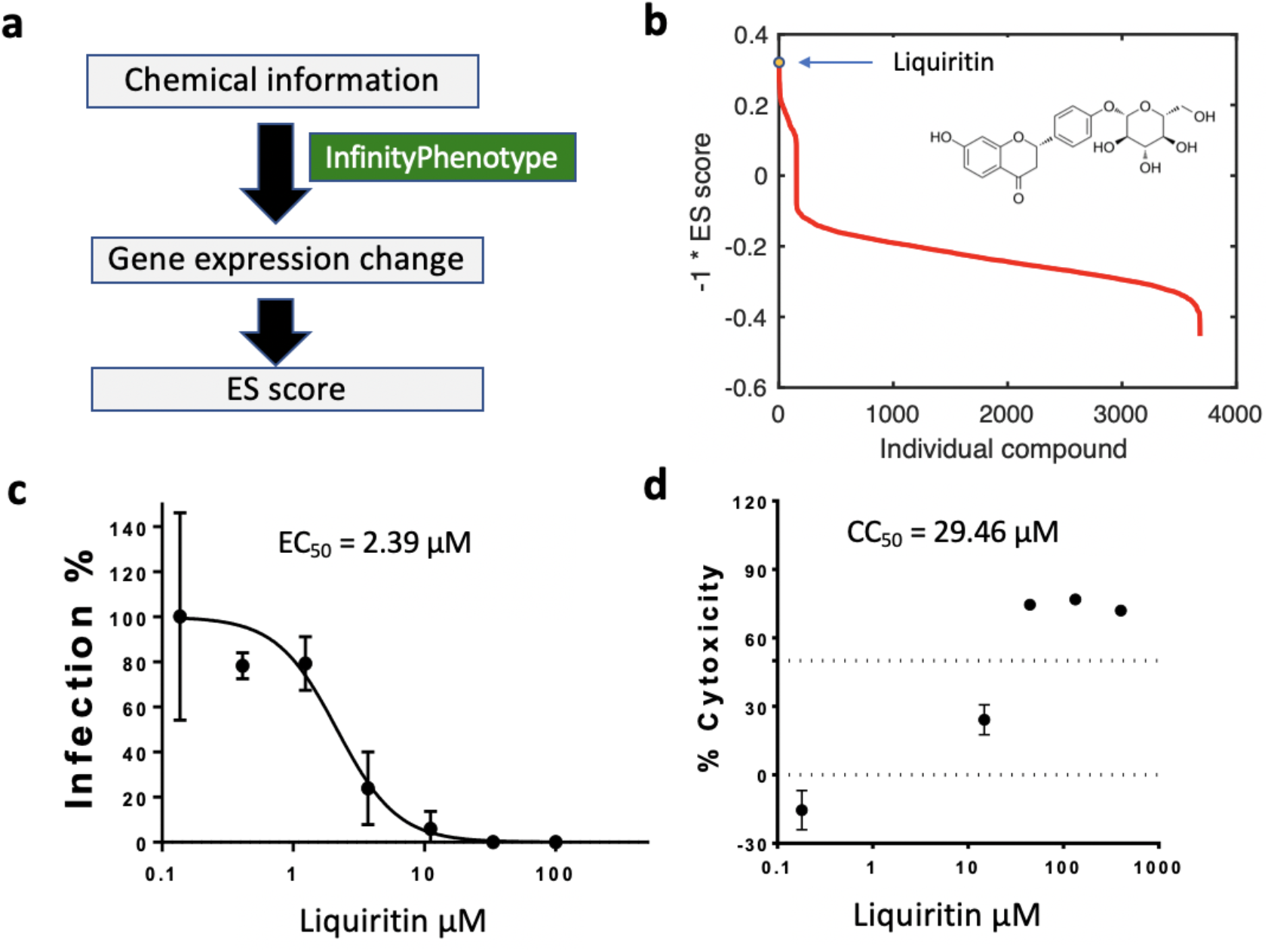
The antiviral activities of the liquiritin against SARS-CoV-2 *in vitro*. a) The workflow of the computation of enrichment score (ES). b) The ES score of all compounds. Liquiritin ranks first. c) The antiviral activity of liquiritin in Vero E6 cells (EC_50_ = 2.39 μM, n = 3 for where error bar is shown). d) Cytotoxicity of the liquiritin to Vero E6 cells was measured by MTS assay.

### In Vitro verification of the efficacy

To experimentally verify the predicted results, we measured the effects of these compounds on the virus yield and infection rates of SARS-CoV-2. First, a cytopathic effect (CPE) inhibition assay with only one dose was performed in Vero E6 cells (ATCC-1586) for fast screening. Among the five tested drugs, liquiritin exhibited significant CPE inhibition at 10 μM qualitatively. We then employed qRT-PCR to quantify the inhibition and expanded the concentration from 0.1 to 100 μM. Vero E6 cells were infected with SARS-CoV-2 at 100 μl median tissue culture infectious dose (TCID_50_). Notably, liquiritin potently blocked virus infection at low-micromolar concentration (EC_50_ = 2.39 μM, Figure 1c).

### Transcriptomic analysis of cell line treated with liquiritin

The transcriptomic change induced by liquiritin (Figure 2a) in MCF7 cells was reported by Li et al^4^. Gene Ontology analysis reveals that the upregulated genes are enriched in the “type I interferon signaling pathway”, “negative regulation of viral genome replication”, “defense response to virus” and “response to virus” categories (Figure 2b) with *p*-value < 8.26e-4, 8.29e-4, 0.0021 and 0.0039, respectively. Nine genes in the first category were IFIT1, IFIT3, IFITM1, IFI6, IFI27, OAS3, OAS1, PSMB8 and IRF8. The induced ISGs include MX1, OAS, IFIT, IFITM and etc. as shown in the “negative regulation of viral genome replication” category. IFIT1 and IFIT3 belong to the type I interferon (IFN)-induced protein with tetratricopeptide repeats (IFIT) family and IFITM1 belongs to the IFN-induced transmembrane protein (IFITM) family. Thus, liquiritin exhibits a general anti-viral activity by mimicking the type I IFN. Further, the enrichment analysis of protein-protein interaction networks reveals a significant module composed of nine ISGs (Figure 2d). Other genes in this module indicate potential regulators of these ISGs.

**Figure 2.**
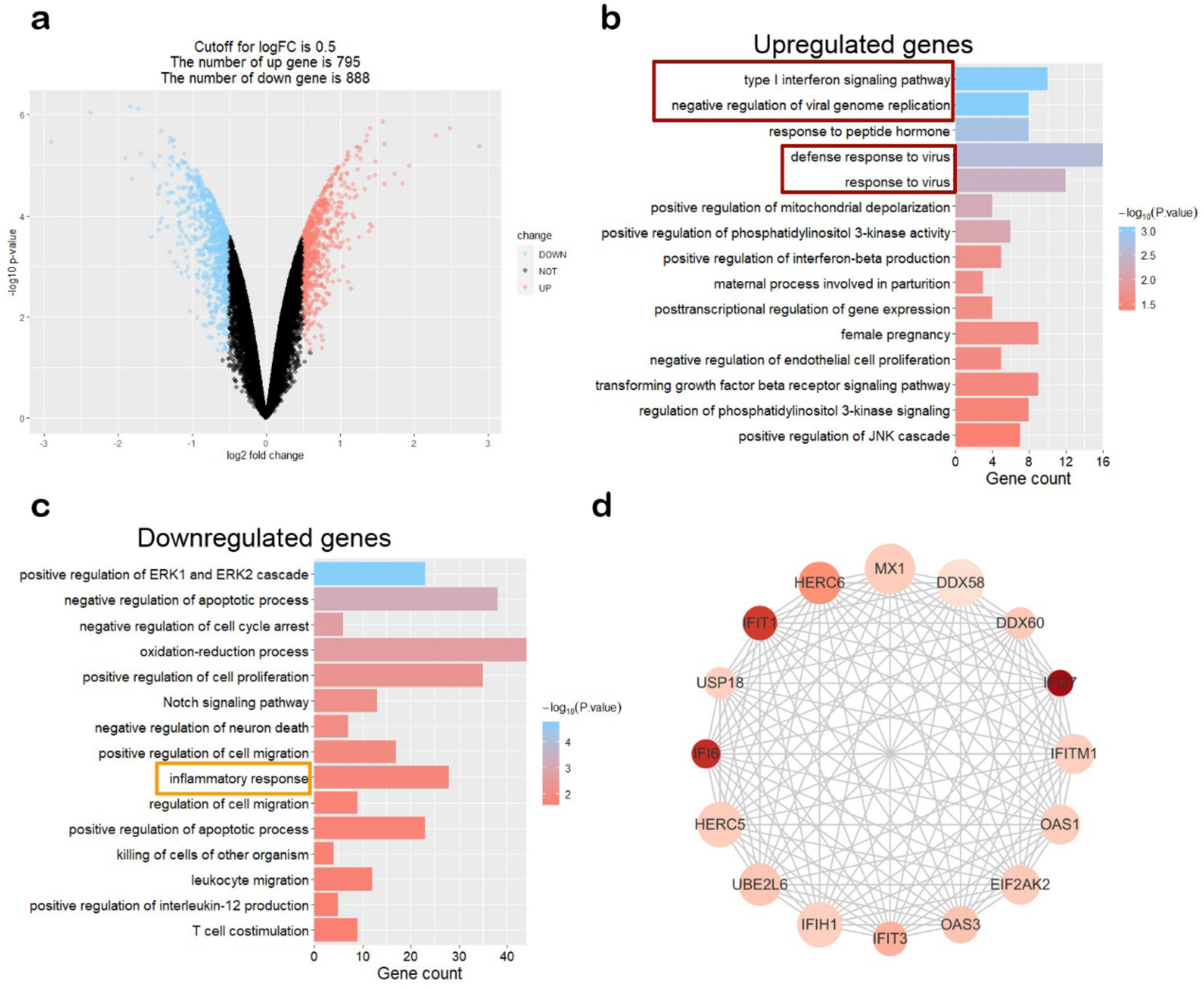
Transcriptomic analysis of cell line treated with liquiritin. a) Volcano plot of the differentially expressed genes (DEGs) in MCF7 cells treated with liquiritin compared to that in normal cells. Gene Ontology analysis of upregulated genes (b) and that of downregulated genes (c). d) The most significant module of the protein-protein interaction (PPI) network of upregulated genes. Node size is positively related to degree of genes and the gradation of color positively associated with the expression change of this gene.

Further, the “inflammatory response” is also enriched in down-regulated genes with *p*-value < 0.015 (Figure 2c), showing liquiritin reduced inflammatory response. In this category, IL13, IL17B and etc. showed up. This may be beneficial for avoiding the cytokine storm observed in COVID-19.

### Liquiritin does not inhibit 3CL^pro^

Liu et al.^18^ found derivatives of flavones efficiently inhibit 3CL^pro^, which is a conserved protease for SARS-CoV-2 replication. Liquiritin is also a derivative of flavones, specifically it is a monosaccharide derivative and a monohydroxyflavanone. It has a mother nucleus structure similar to the compounds found by Liu et al. Though molecule docking showed there might be a binding to 3CL^pro^, liquiritin didn’t show any inhibitory effects to the activity of 3CL^pro^ *in vitro* at 12.5, 25.0 and 50.0 μM (data not shown).

### Liquiritin is a safe candidate for treating COVID-19

To evaluate the safety of liquiritin, the cytotoxicity in Vero E6 cells was determined by MTS assay (CC_50_ = 29.46 μM, Maximal inhibition of cell growth is 70%, as shown in Figure 1d). We further examined the toxicity in vivo. No morality was observed at 150 mg/kg dose of one-time treatment of liquiritin by intravenous administration during the 14 days monitoring period (Figure 3a). Mice intravenous administration 150 mg/kg of liquiritin did not manifest any signs of toxicity during this observation period. The body weight, food intake and water intake are shown in Figure 3b, c. A continuous treatment for seven days at 300 mg/kg was also performed. The mortality rate is also zero (Figure 3d). The intake food, body weights were all kept at normal level (Figure 3e, f). As reported in literatures, liquiritin has no significantly cytotoxic effects in L-02, GES-1, IMR-90 and 293T cells with different concentrations (30, 60, 90, 120, and 150 μM) and the MTT assay results confirmed that 345 μM liquiritin has no influence on the viability of RA-FLS cells^16,17^.

**Figure 3.**
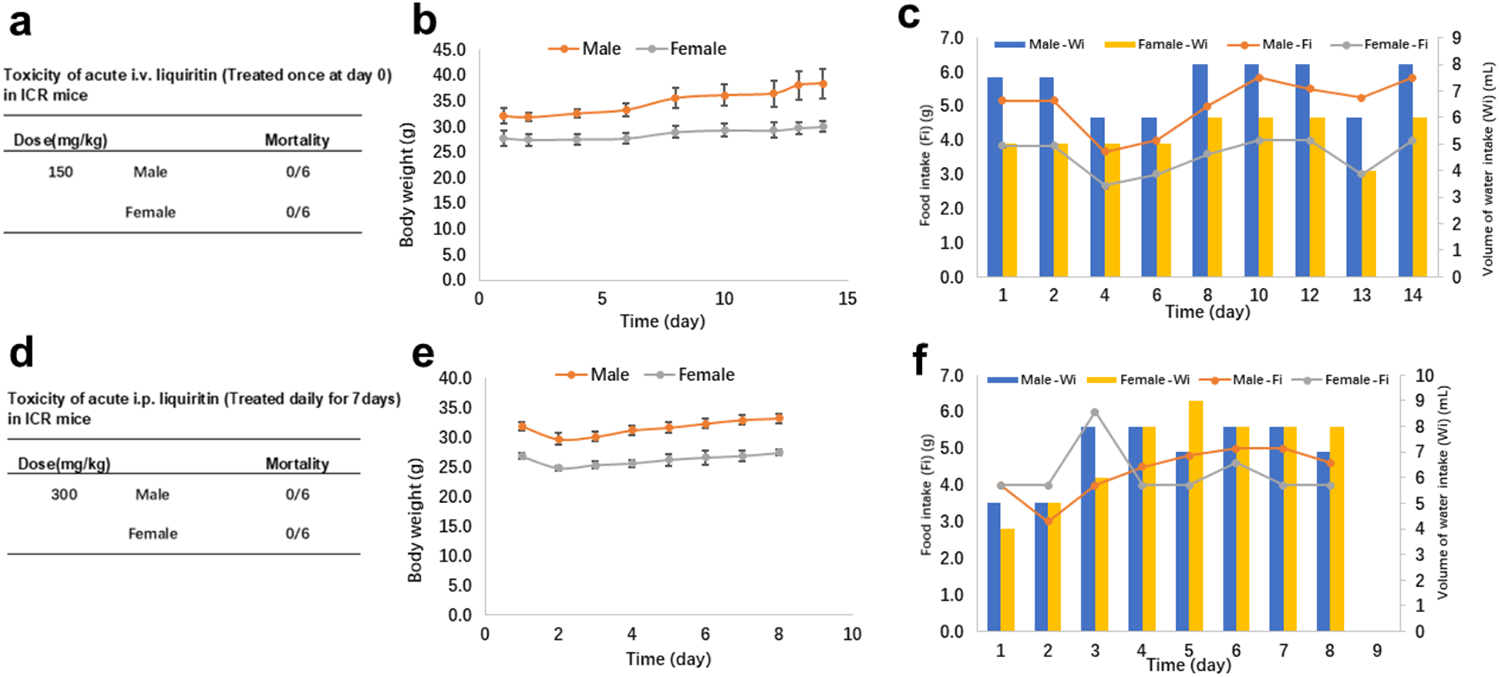
The Toxicity assay for liquiritin in ICR mice. a) Mortality rate of acute i.v. liquiritin in ICR mice. b, c) Body weight, food intake and water intake at 150 mg/kg dose of one-time treatment of liquiritin by intravenous administration during the 14 days monitoring period in ICR mice. d) Mortality rate of acute i.p. liquiritin in ICR mice. e, f) Body weight, food intake and water intake at 300 mg/kg dose of liquiritin by i.p. administration for seven days in ICR mice.

For both assays shown in Figure 3a, d, we also measured organ and serum biochemical indexes. No significant differences organ indexes (ratio of organ weight to body weight) were observed except liver index compared with the control mice. Meanwhile, levels of glucose, cholesterol, triglyceride, high-density lipoprotein (HDL-C), and low-density lipoprotein (LDL-C) kept normal after liquiritin administration. We observed declined ALT and it may be due to the anti-inflammatory effect of liquiritin. Meanwhile, no abnormality was observed in levels of blood UREA. UA. CK. α-HBDH, suggesting renal function and heart function were not harmed. Though some relevant index of blood analysis, such as the number of leukocytes, neutrophils and etc., in the control mice and liquiritin-treated mice are different, they are all within the range of reference values, indicating that liquiritin has no obvious toxicity (Supplementary Table 6–9).

### Pharmacokinetics survey of liquiritin

To evaluate the pharmacokinetics of liquiritin, we searched literatures and found the pharmacokinetic data of mixtures containing liquiritin (Supp. Table 2). The corresponding dose of liquiritin is range from 225 to 44, 000 ug/kg, the C_max_ in serum is range from 0.064 to 2.3 μM. The T_max_ is range from 0. 25 to 0. 6 hours and T_1/2_ is range from 1.9 to 11.63 hours. As the bioavailability is low and degradation is fast, we suggest an intravenous or inhalation administration for liquirtin.

## Discussion

To find drugs that may cure COVID-19, we adopted a different strategy by analyzing transcriptional changes induced by various compounds. We found that liquiritin is highly effective in the inhibition of SARS-CoV-2 infection *in vitro* compared to other SARS-CoV-2 drugs reported *in vitro* (Supp. Table 3). Since liquiritin is also one of the main ingredients of compound licorice tablets, we speculate that it is safe. Indeed, it didn’t show toxicity and side effects in two independent experiments. In summary, we suggest that liquiritin should be assessed in human patients suffering from the COVID-19.

Mechanistically, we found liquiritin induced the expression of anti-viral genes. GO analysis revealed that “type I INF signaling pathway” was activated. We thus proposed that liquiritin may mimic type I INF to inhibit the viral replication. Vero E6 cell line we used here lack IFNB1 gene and cannot synthesize interferon. However, Vero E6 cells contain the *trans*-acting factors needed for human beta interferon^20^. It’s still responsive to human INF. Thus, liquiritin acted through downstream of INF synthesis.

We also surveyed the literatures for the targets of liquiritin. It’s reported that liquiritin, a flavonoid isolated from Glycyrrhiza, is a potent and competitive AKR1C1 inhibitor with IC_50_ of 0.62 μM, 0.61 μM, and 3.72μM for AKR1C1, AKR1C2 and AKR1C3, respectively^19^. Whether AKR1 is the major target for liquiritin’s anti-viral activity is unclear. Further experiments need to be carried out.

So far, there are no first principles to precisely simulate the intrinsic interactions of cells. One alternative solution is to use the measured data to overcome the difficulty of complex systems. Deep neural network offers an option to empirically fit high dimensional data. InfinityPhenotype we employed here is based on transcriptional change induced by compounds. Such framework treated a broad range of interactions inside the cells, from ligand-protein interaction to transcriptional regulations, as well as other hidden interactions, as a black box. It offers us an opportunity to predict the efficacy of compounds in a precise manner.

To test if other gradients in liquorice inhibit the viral replication, we also tested liquiritigenin and glycyrrhizic acid in Vero E6 cells. Neither of them showed inhibitory effect (data not shown). Further animal test, transcriptional measurement with virus in Vero E6 cells and in mice are still ongoing. Because type I interferon is responsive to most viruses that infect vertebrate, we expect liquiritin having a broad and potent antiviral function to other viral pathogens, like HBV, HIV and etc.

## Methods

### Virus and Cells

The stock of SARS-CoV-2 strain BetaCoV/wuhan/AMMS01/2020 was originally isolated from a patient returning from Wuhan. The virus was amplified and titerated by standard plaque forming assay on Vero cells. All experiments involving infectious SARS-CoV-2 were performed in biosafety level 3 (BSL3) containment laboratory in AMMS. Vero cell were purchased from ATCC (Cat# CCL-81) and cultured in DMEM supplemented with 10% FBS at 37%.

### Prediction of efficacy

The full list of genes used for enrichment analysis (GSEA) includes SLC1A5, CXADR, CAV2, NUP98, CTBP2, GSN, HSPA1B, STOM, RAB1B, HACD3, ITGB6, IST1, NUCKS1, TRIM27, APOE, SMARCB1, UBP1, CHMP1A, NUP160, HSPA8, DAG1, STAU1, ICAM1, CHMP5, DEK, VPS37B, EGFR, CCNK, PPIA, IFITM3, PPIB, TMPRSS2, UBC, LAMP1 and CHMP3. Kolmogorov-Smirnov (KS) test is used to evaluate the distribution of query genes in reference list. The gene expression fold change was calculated by InfinityPhenotype platform and used to make the rank list (>10,000 in this case). The ES score for one gene set is defined as:

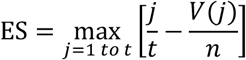

where t is the number of genes in the targeted gene set, n is the number of genes in full gene expression profile, V(j) is the rank of a specific gene in the rank list.

### Cytotoxicity assay

The cytotoxicity of the tested drugs on Vero Cell were determined by MTS cell proliferation assays (Promega, USA), according to manufacturer’s protocol. Briefly, ~1 × 104 cells were seeded into a 96-well plate and incubated for 20-24 h at 37 °C. The media was removed, and 100 μl of media containing decreasing concentrations of antiviral compound were added to the wells. All determinations were performed in triplicate. After 4 days incubation at 37 °C, MTS assays were performed according to manufacturers’ protocols. A microtiter plate reader (SYNERGY HTX, BioTek Instruments) with a 490 nm filter was used to record absorbance. After adjusting the absorbance for background and comparing to untreated controls, the cytotoxic concentration CC50 was calculated using a sigmoidal nonlinear regression function to fit the dose-response curve using the GraphPad Prism 7.0 software.

### The in vitro antiviral activity

The in vitro antiviral efficacy of the tested drug on Vero E6 Cell was performed as we previously described [Jin, et al., 2020]. Briefly, ~1 × 104 cells were seeded into a 96-well plate and incubated for 20-24 h at 37 °C. Cells were pre-treated with the indicated antivirals (10 μM) for 1 h, and the virus (MOI of 0.01) was subsequently added to allow infection for 2 h. Then, the virus-drug mixture was removed and cells were further cultured with fresh drug-containing medium. At 48 h p.i., viral RNA copy in supernatants were quantified by qRT-PCR assays with the SARS-CoV-2-specific primers.

### Toxicity study in ICR mice

ICR mice were used to test the in vivo safety of liquiritin. Thirty-Six ICR male and female mice were purchased from the Peking University Animal Department. The animals were housed in a temperature-controlled animal room (24 ± 2 °C) with a relative humidity of 60-80%. All mice were fasted overnight but given water ad libitum prior to dosage. Animals were divided into two groups with 6 males and 6 females at random. Liquiritin was dissolved in 10% DMSO/40% PEG400/50% Saline and administered i.v. once at doses of 150mg/kg, and administered i.p. once a day for 10 days at doses of 300mg/kg. Their general behavior, signs of toxicity, body weights and mortality were recorded after the administration of liquiritin. Animals were killed by excessive anesthesia. Gross necropsy, weighing, calculation of organ index (brain, heart, lung, liver, kidney, adrenal gland, thymus, spleen, testis, epididymis, ovary, uterus, thyroid), pathological examination of the organs was carried out afterwards. The blood sample was analysis by blood analyzer HEMAVET 950FS.

### Transcriptional analysis

Microarray data on gene expression (GSE85871) was downloaded from Gene Expression Omnibus (GEO). The DEGs between normal and liquirintin-treated MCF7 cells were also screened by the limma package. |log2FC| ≥ 0.5, adjust p-value < 0.05 were considered statistically significant for the DEGs. To elucidate potential biological process associated with the DEGs, we performed GO enrichment analysis utilizing the Database for Annotation, Visualization and Integrated Discovery (DAVID). p-value < 0.05 was defined as the cut-off criteria. The upregulated DEGs were mapped to STRING to evaluate the PPI information and set confidence score > 0.4 as the cut-off standard. Cytoscape was used to visualize the PPI network, a practical open–source software tool for the visual exploration of biomolecule interaction networks consisting of protein, gene and other types of interaction.

### Enzyme Inhibition Assay

A colorimetric substrate Thr-Ser-Ala-Val-Leu-Gln-pNA (GL Biochemistry Ltd) and assay buffer (40 mM PBS, 100 mM NaCl, 1 mM EDTA, 0.1% Triton 100, pH 7.3) was used for the inhibition assay. Stock solutions of the inhibitor were prepared with 100% DMSO. The 100 μl reaction systems in assay buffer contain 0.5 μM protease and 5% DMSO or inhibitor to the final concentration. First, dilute SARS-CoV-2 3CLpro with assay buffer to the desired concentration. 5 μl DMSO or inhibitor at various concentrations was pre-incubated with 85 μl diluted SARS-CoV-2 3CLpro for 30 min at room temperature. And then add 10 μl 2 mM substrate Thr-Ser-Ala-Val-Leu-Gln-pNA (dissolved in water) into above system to final concentration of 200 μM to initiate the reaction. Increase in absorbance at 390 nm was recorded for 20 min at interval of 30 s with a kinetics mode program using a plate reader (Synergy, Biotek). The percent of inhibition was calculated by V_i_/V_0_, where V_0_ and V_i_ represent the mean reaction rate of the enzyme incubated with DMSO or compounds. IC_50_ was fitted with Hill function.

## Author Contributions

All authors contributed to the work presented in this paper. J. Z. & W. K. performed the statistical analysis, Z. L. and Z. X performed the efficacy prediction, Y. D., X. L, and N. Z measured the anti-viral activity and performed the analysis. B. Z. performed docking, J. Z. performed the literature review. Z. X & J. Z. wrote the paper. Z. X & C. Q. supervised the team.

## Acknowledgement

We thank Hongbo Liu and Luhai Lai for the help of 3CL^pro^ assay. Z.W.X was supported by National key research and development program of China (2018YFA0900200), National Natural Science Foundation of China Grants (31771519) and Beijing Natural Science Foundation (5182012). C.F.Q. was supported by the National Science Fund for Distinguished Young Scholar (No. 81925025), and the Innovative Research Group (No. 81621005) from the NSFC, and the Innovation Fund for Medical Sciences (No.2019-I2M-5-049) from the Chinese Academy of Medical Sciences.

---

**Supplementary Table 1.** Results of InfinityPhenotype platform screening

| ID | Name | Drug Groups | score |
| --- | --- | --- | --- |
| T7S0344 | Liquiritin | natural<br>product | -0.315809062 |
| T3S1363 | Agnuside | natural<br>product | -0.308751956 |
| DB01366 | Procaterol | approved;<br>investigational | -0.233934229 |
| DB13878 | Pibrentasvir | approved;<br>investigational | -0.2277026 |
| DB04339 | Carbocisteine | approved;<br>investigational | -0.212099668 |

**Supplementary Table 2.** Pharmacokinetic Data of Mixtures Containing Liquiritin

| Animal | Compounds | Liquiritin dose | Route | Pharmacokinetic parameters | Ref |
| --- | --- | --- | --- | --- | --- |
|  | Glycyrrhizae |  |  | C<sub>max</sub> : 0.064μM |  |
| Rat | Radix | 3.8mg/kg | oral | T<sub>max</sub> : 0.4h | (1) |
|  | Extract |  |  | T<sub>1/2</sub> : 3.7h |  |
|  | Chaihu- |  |  | C<sub>max</sub> : 0.5μM |  |
| Rat | Guizhi | 225μg/kg | oral | T<sub>max</sub> : 0.11h | (2) |
|  | decoction |  |  | T<sub>1/2</sub> : 11.63h |  |
|  | Tongmai |  |  | C<sub>max</sub> : 0.475μM |  |
| Rat | Yangxin Pill | 1.337mg/kg | oral | T<sub>max</sub> : 0.5h | (3) |
|  |  |  |  | T<sub>1/2</sub> : 2.16h |  |
|  | ZBPYR |  |  | C<sub>max</sub> : 0.5μM |  |
| Rat | extract | 6.25mg/kg | oral | T<sub>max</sub> : 0.6h | (4) |
|  |  |  |  | T<sub>1/2</sub> : 7.8h |  |
| Rat | Baoyuan | 240.17/600.425 | oral | C<sub>max</sub> : 0.077/ | (5) |
|  | Decoction | /1200μg/kg |  | 0.1156/0.14μM |  |
|  | Si-Ni-San |  |  | C<sub>max</sub> : 2.3μM |  |
| Rat | extract | 16.76mg/kg | oral | T<sub>max</sub> : 0.5h | (6) |
|  |  |  |  | T<sub>1/2</sub> : 7.25h |  |
|  | Longhu |  |  | C<sub>max</sub> : 0.24μM |  |
| Rat | Rendan pills | 2.07mg/kg | oral | T<sub>max</sub> : 0.3h | (7) |
|  |  |  |  | T<sub>1/2</sub> : 1.9h |  |
|  | Ge-Gen |  |  | C<sub>max</sub> : 0.41μM |  |
| Rat | Decoction | 44.0mg/kg | oral | T<sub>max</sub> : 0.25h | (8) |
| (female) |  |  |  | T<sub>1/2</sub> : 4.26h |  |

**Supplementary Table 3.**
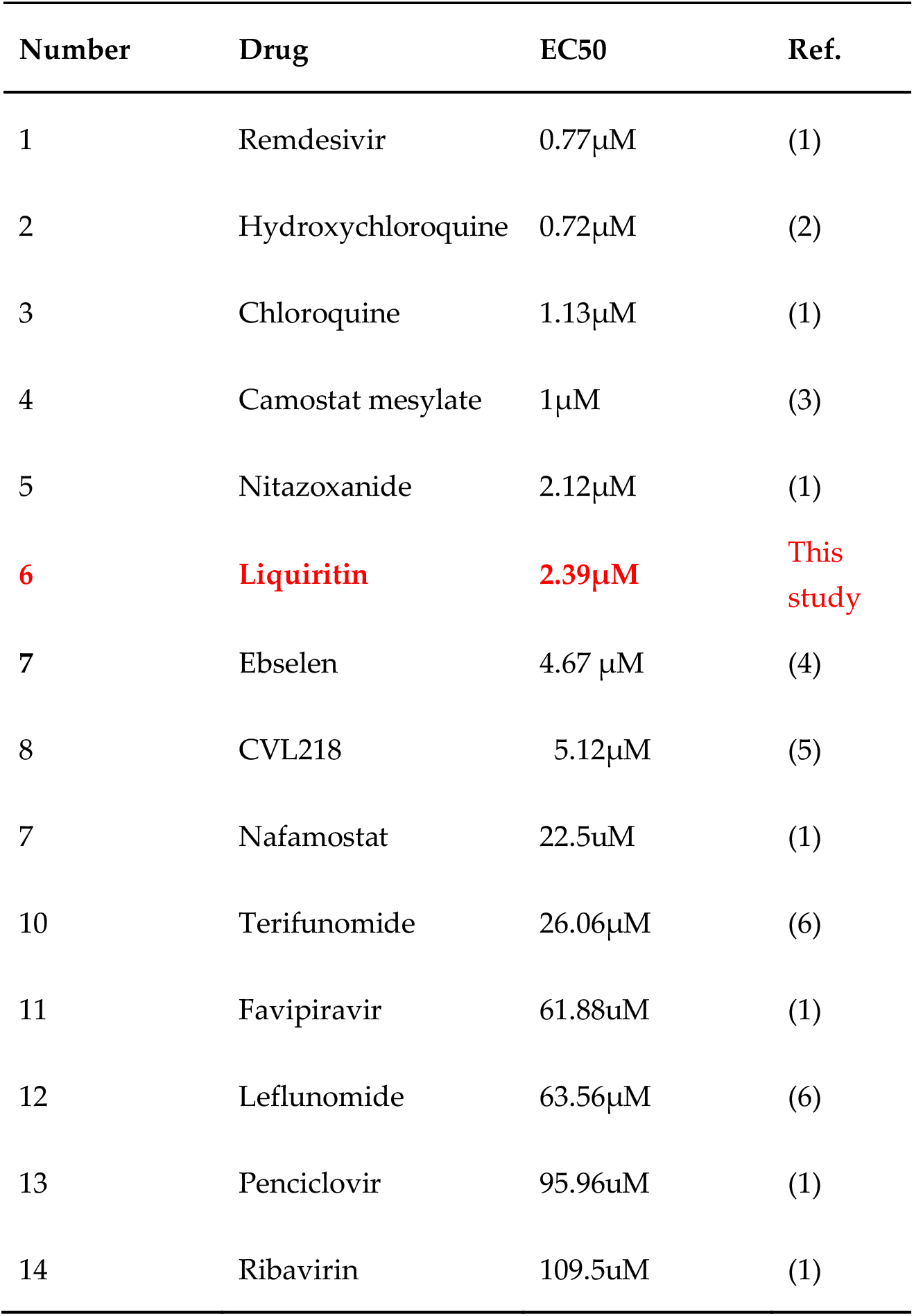
Against SARS-CoV-2 drugs reported *in vitro* and Liquiritin

**Supplementary Table 4.**
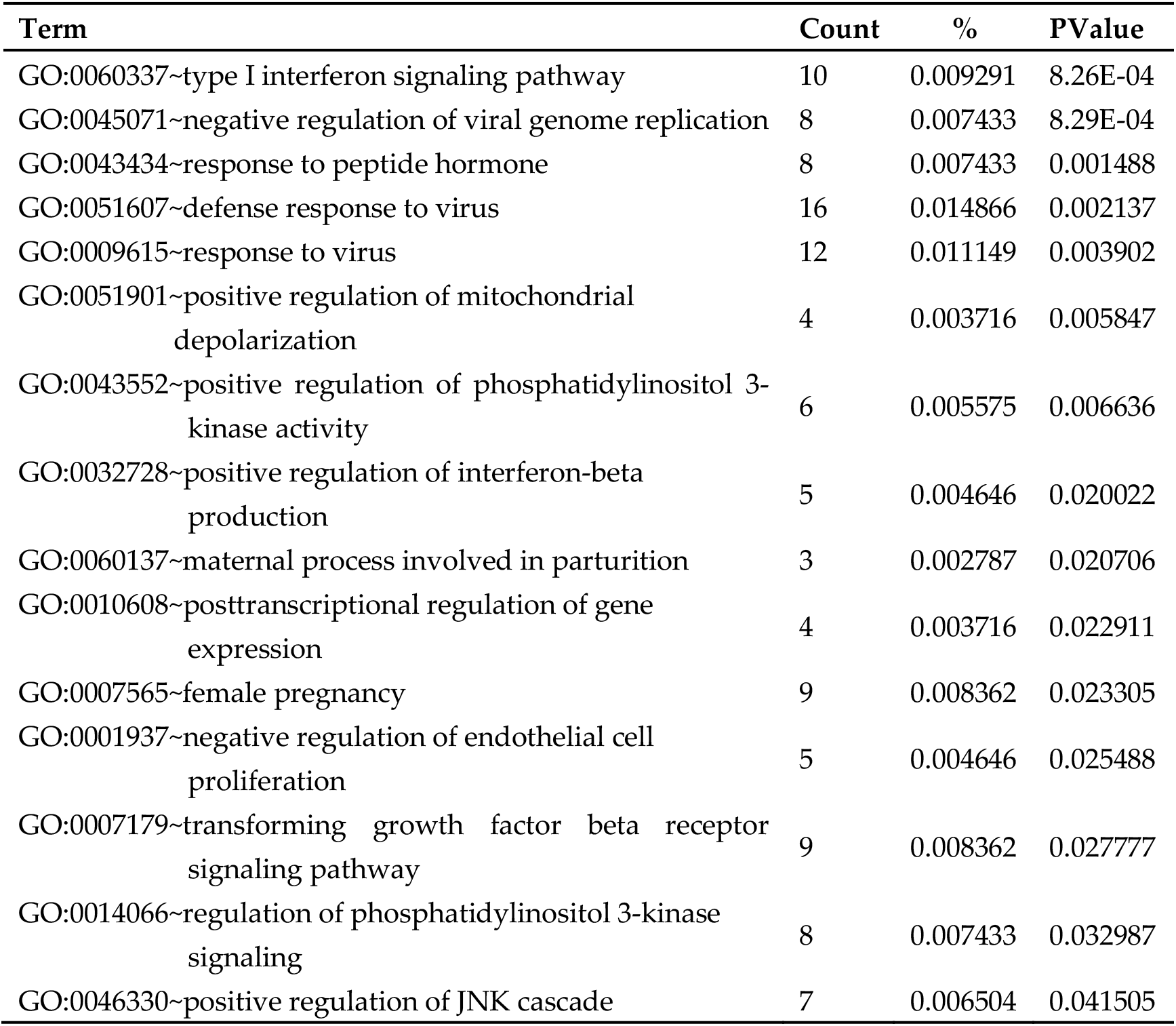
GO analysis of upregulated genes

**Supplementary Table 5:** GO analysis of downregulated genes

| **Term** | **Count** | **%** | **PValue** |
| --- | --- | --- | --- |
| GO:0070374~positive regulation of ERK1 and ERK2 cascade | 23 | 0.019502 | 1.65E-05 |
| GO:0043066~negative regulation of apoptotic process | 38 | 0.032221 | 5.22E-04 |
| GO:0071157~negative regulation of cell cycle arrest | 6 | 0.005087 | 0.001725 |
| GO:0055114~oxidation-reduction process | 44 | 0.037308 | 0.00186 |
| GO:0008284~positive regulation of cell proliferation | 35 | 0.029677 | 0.004883 |
| GO:0007219~Notch signaling pathway | 13 | 0.011023 | 0.006354 |
| GO:1901215~negative regulation of neuron death | 7 | 0.005935 | 0.009122 |
| GO:0030335~positive regulation of cell migration | 17 | 0.014414 | 0.010436 |
| GO:0006954~inflammatory response | 28 | 0.023741 | 0.015459 |
| GO:0030334~regulation of cell migration | 9 | 0.007631 | 0.019467 |
| GO:0043065~positive regulation of apoptotic process | 23 | 0.019502 | 0.020164 |
| GO:0031640~killing of cells of other organism | 4 | 0.003392 | 0.023776 |
| GO:0050900~leukocyte migration | 12 | 0.010175 | 0.024075 |
| GO:0032735~positive regulation of interleukin-12 production | 5 | 0.00424 | 0.025538 |
| GO:0031295~T cell costimulation | 9 | 0.007631 | 0.025874 |

**Supplementary Table 6:**
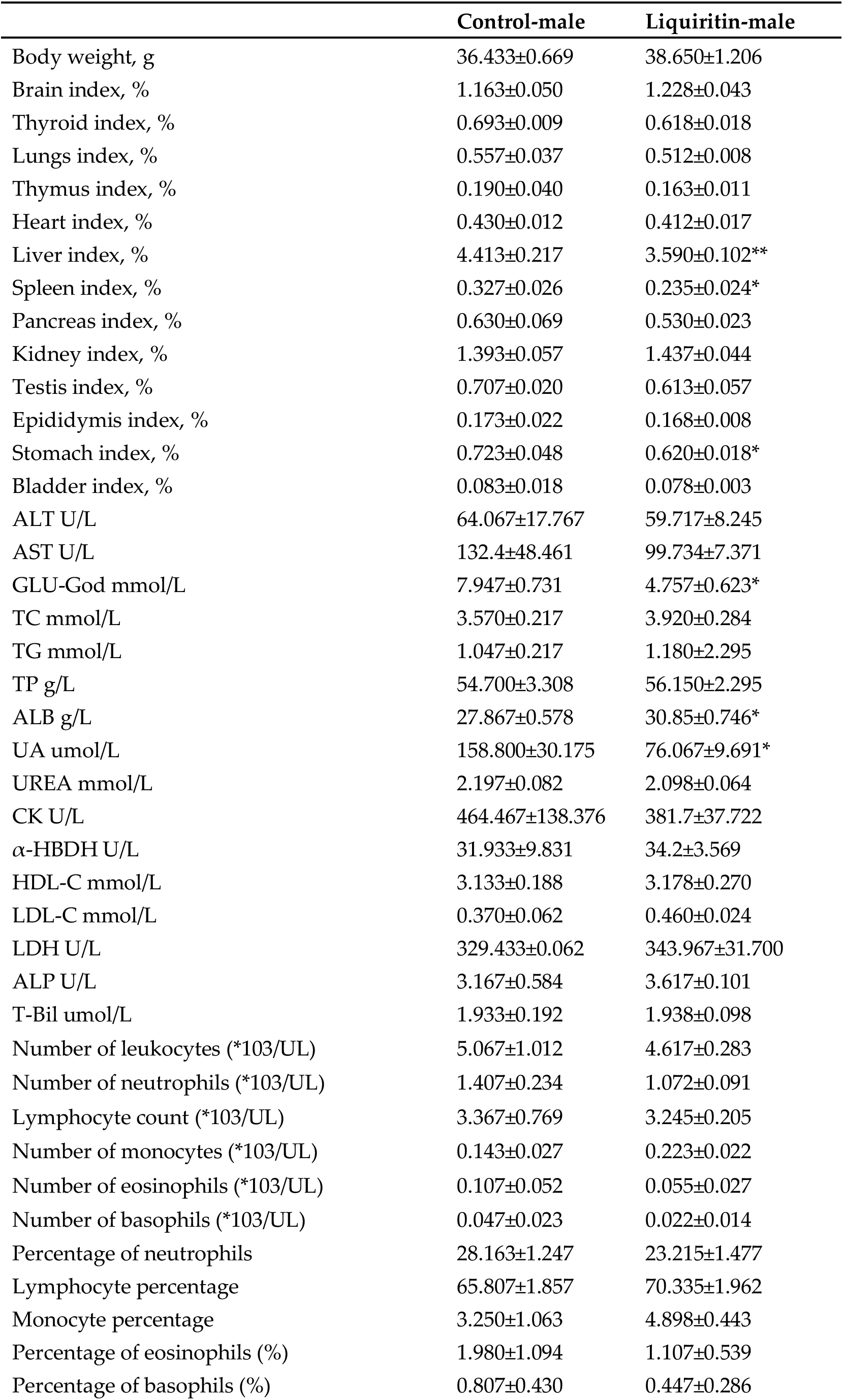
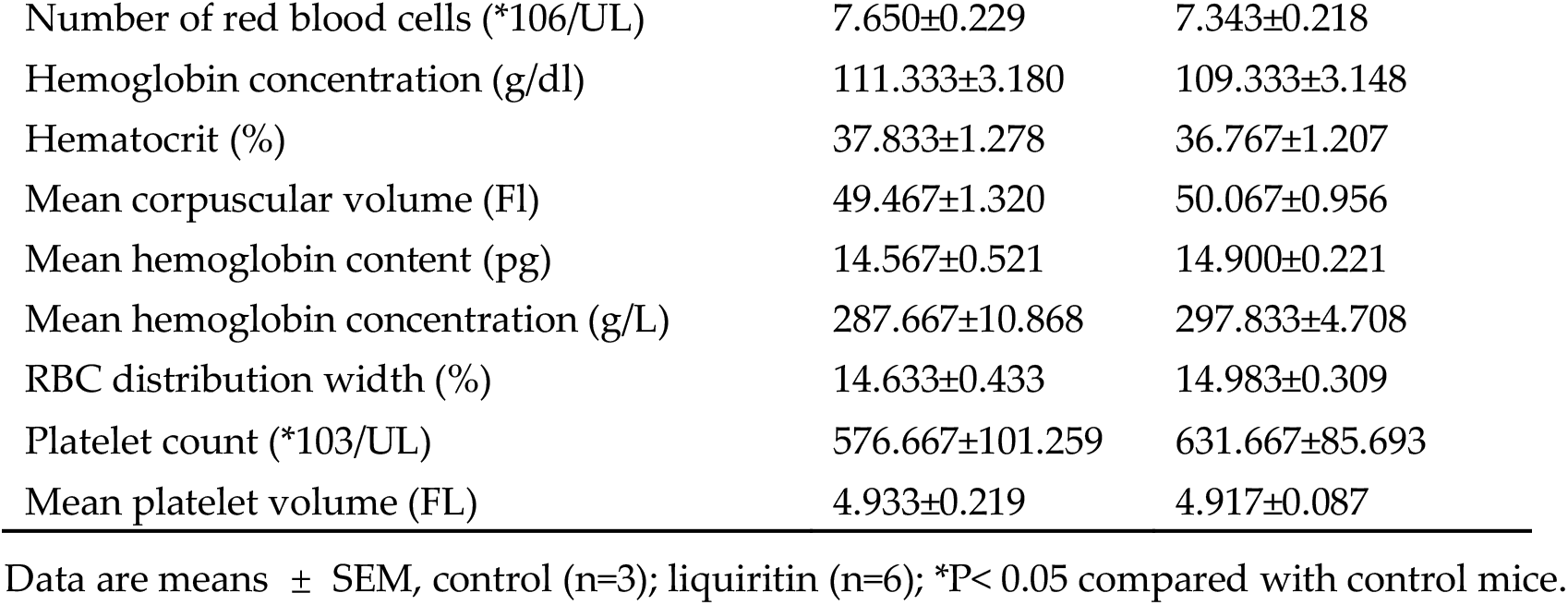
Body weight, organ index, blood chemistry and blood analysis in control or liquiritin-treated male mice by intravenous injection.

**Supplementary Table 7:** Body weight, organ index, blood chemistry and blood analysis in control or liquiritin-treated female mice by intravenous injection.

|  | Control-female | Liquiritin-female |
| --- | --- | --- |
| Body weight, g | 28.867±0.581 | 30.283±0.473 |
| Brain index, % | 1.583±0.041 | 1.520±0.035 |
| Thyroid index, % | 0.647±0.034 | 0.680±0.042 |
| Lungs index, % | 0.590±0.050 | 0.612±0.013 |
| Thymus index, % | 0.183±0.022 | 0.215±0.018 |
| Heart index, % | 0.427±0.009 | 0.428±0.018 |
| Liver index, % | 4.393±0.170 | 3.688±0.054** |
| Spleen index, % | 0.403±0.015 | 0.363±0.021 |
| Pancreas index, % | 0.600±0.036 | 0.597±0.024 |
| Kidney index, % | 1.200±0.055 | 1.145±0.037 |
| Ovary index, % | 0.090±0.010 | 0.127±0.010 |
| Uterus index, % | 0.430±0.075 | 0.520±0.065 |
| Stomach index, % | 0.817±0.048 | 0.722±0.032 |
| Bladder index, % | 0.070±0.000 | 0.065±0.002 |
| ALT U/L | 56.133±5.441 | 40.400±2.912* |
| AST U/L | 107.4±12.450 | 118.350±2.949 |
| GLU-God mmol/L | 5.873±0.779 | 5.315±0.772 |
| TC mmol/L | 2.65±0.560 | 2.968±0.166 |
| TG mmol/L | 0.617±0.116 | 0.740±0.080 |
| TP g/L | 55.333±3.290 | 61.083±1.554 |
| ALB g/L | 30.600±0.321 | 34.183±0.715* |
| UA umol/L | 141.4±56.486 | 58.583±2.483 |
| UREA mmol/L | 2.757±0.472 | 1.998±0.049* |
| CK U/L | 442.267±38.585 | 439.733±26.190 |
| α-HBDH U/L | 28.633±1.962 | 32.383±3.144 |
| HDL-C mmol/L | 2.317±0.462 | 2.627±0.122 |
| LDL-C mmol/L | 0.410±0.076 | 0.287±0.034 |
| LDH U/L | 290.933±38.479 | 325.017±20.649 |
| ALP U/L | 2.467±0.674 | 3.533±0.314 |
| T-Bil umol/L | 1.160±0.096 | 1.308±0.062 |
| Number of leukocytes (*103/UL) | 5.327±0.280 | 3.837±0.686 |
| Number of neutrophils (*103/UL) | 1.353±0.246 | 0.937±0.229 |
| Lymphocyte count (*103/UL) | 3.670±0.156 | 2.640±0.432 |
| Number of monocytes (*103/UL) | 0.197±0.048 | 0.188±0.023 |
| Number of eosinophils (*103/UL) | 0.080±0.056 | 0.052±0.027 |
| Number of basophils (*103/UL) | 0.027±0.018 | 0.017±0.012 |
| Percentage of neutrophils | 25.150±3.512 | 22.780±2.326 |
| Lymphocyte percentage | 69.340±5.048 | 69.998±1.521 |
| Monocyte percentage | 3.623±0.667 | 5.710±1.173 |
| Percentage of eosinophils (%) | 1.443±0.931 | 1.092±0.568 |
| Percentage of basophils (%) | 0.447±0.294 | 0.422±0.259 |
| Number of red blood cells (*10<sup>6</sup>/UL) | 7.123±0.138 | 7.485±0.262 |
| Hemoglobin concentration (g/dl) | 109.333±4.055 | 107.833±2.713 |
| Hematocrit (%) | 39.100±0.577 | 35.900±1.309 |
| Mean corpuscular volume (fL) | 54.933±0.348 | 48.050±1.447 |
| Mean hemoglobin content (pg) | 15.200±0.404 | 14.450±0.339 |
| Mean hemoglobin concentration (g/L) | 277.000±6.245 | 301.500±7.261 |
| RBC distribution width (%) | 14.867±0.467 | 15.733±0.226 |
| Platelet count (*10<sup>3</sup>/UL) | 532.667±42.881 | 536.333±66.231 |
| Mean platelet volume (fL) | 5.100±0.153 | 5.050±0.099 |

Data are means ± SEM, control (n=3); liquiritin (n=6); \*P< 0.05 compared with control mice.

**Supplementary Table 8:** Body weight, organ index, blood chemistry and blood analysis in control or liquiritin-treated male mice by intraperitoneal injection.

|  | Control-male | Liquiritin-male | Liquiritin-male-recovery |
| --- | --- | --- | --- |
| Body weight, g | 35.133±0.426 | 34.533±0.240 | 34.600±0.321 |
| Brain index, % | 1.243±0.038 | 1.307±0.024 | 1.320±0.046 |
| Thyroid index, % | 0.770±0.050 | 0.703±0.038** | 0.787±0.048 |
| Lungs index, % | 0.537±0.012 | 0.570±0.040 | 0.540±0.035 |
| Thymus index, % | 0.150±0.010 | 0.220±0.010* | 0.173±0.003 |
| Heart index, % | 0.420±0.015 | 0.387±0.012 | 0.443±0.013 |
| Liver index, % | 4.607±0.239 | 3.907±0.058 | 5.847±0.116* |
| Spleen index, % | 0.267±0.027 | 0.417±0.035 | 0.343±0.018 |
| Pancreas index, % | 0.567±0.055 | 0.610±0.025 | 0.817±0.067 |
| Kidney index, % | 1.687±0.058 | 1.437±0.047 | 1.427±0.084 |
| Testis index, % | 0.683±0.009 | 0.617±0.029 | 0.663±0.009 |
| Epididymis index, % | 0.143±0.003 | 0.143±0.018 | 0.183±0.024 |
| Stomach index, % | 0.737±0.050 | 0.657±0.022 | 0.670±0.038 |
| Bladder index, % | 0.083±0.015 | 0.080±0.010 | 0.080±0.010 |
| ALT U/L | 131.833±27.310 | 45.100±5.856* | 57.833±6.743 |
| AST U/L | 137.000±3.555 | 113.100±14.755 | 118.767±10.810 |
| GLU-God mmol/L | 3.753±0.387 | 3.680±0.996 | 9.350±0.356*** |
| TC mmol/L | 2.970±0.119 | 3.750±0.545 | 3.320±0.118 |
| TG mmol/L | 0.573±0.047 | 0.940±0.126 | 1.227±0.172* |
| TP g/L | 71.800±19.310 | 60.100±3.600 | 55.133±0.617 |
| ALB g/L | 27.167±1.097 | 31.767±1.330 | 28.233±0.524 |
| UA umol/L | 74.967±4.101 | 106.567±16.136 | 53.900±1.504** |
| UREA mmol/L | 2.217±0.119 | 1.933±0.027 | 2.237±0.182 |
| CK U/L | 335.200±72.660 | 481.600±53.306 | 517.767±25.695 |
| α-HBDH U/L | 39.900±5.900 | 34.433±9.089 | 30.067±4.784 |
| HDL-C mmol/L | 2.383±0.084 | 2.900±0.454 | 2.887±0.061** |
| LDL-C mmol/L | 0.477±0.033 | 0.613±0.062 | 0.360±0.066 |
| LDH U/L | 443.900±48.916 | 433.600±112.348 | 314.100±54.673 |
| ALP U/L | 3.267±0.333 | 2.567±0.318 | 5.267±0.145** |
| T-Bil umol/L | 1.533±0.194 | 1.310±0.140 | 1.760±0.068 |
| Number of leukocytes(*103/UL) | 6.020±0.652 | 5.700±0.495 | 7.513±0.615 |
| Number of neutrophils(*103/UL) | 1.600±0.186 | 2.907±0.257* | 1.607±0.279 |
| Lymphocyte count(*103/UL) | 3.890±0.586 | 2.190±0.262 | 5.543±0.312 |
| Number of monocytes(*103/UL) | 0.337±0.041 | 0.547±0.166 | 0.020±0.006** |
| Number of eosinophils(*103/UL) | 0.140±0.015 | 0.050±0.010** | 0.340±0.080 |
| Number of basophils(*103/UL) | 0.047±0.020 | 0.003±0.003 | 0.003±0.003 |
| Percentage of neutrophils | 27.093±4.170 | 51.470±5.110* | 21.063±2.056 |
| Lymphocyte percentage | 64.043±4.323 | 38.310±2.880** | 74.157±2.703 |
| Monocyte percentage | 5.613±0.221 | 9.257±2.361 | 0.250±0.104*** |
| Percentage of eosinophils(%) | 2.383±0.377 | 0.863±0.127* | 4.500±0.761 |
| Percentage of basophils (%) | 0.870±0.090 | 0.103±0.055** | 0.027±0.018*** |
| Number of red blood cells (*106/UL) | 7.437±0.383 | 7.790±0.145 | 7.967±0.072 |
| Hemoglobin concentration (g/dl) | 104.000±4.041 | 114.000±1.000 | 111.667±3.180 |
| Hematocrit (%) | 36.533±1.081 | 38.767±1.444 | 35.367±0.581 |
| Mean corpuscular volume (fL) | 49.300±1.909 | 49.767±0.949 | 44.367±0.437 |
| Mean hemoglobin content (pg) | 14.000±0.404 | 14.633±0.133 | 14.033±0.273 |
| Mean hemoglobin concentration(g/L) | 284.333±3.930 | 294.667±8.743 | 315.333±3.756** |
| RBC distribution width (%) | 15.533±0.426 | 16.967±0.145* | 16.567±0.780 |
| Platelet count (*103/UL) | 438.333±90.381 | 737.000±24.880* | 771.333±33.418* |
| Mean platelet volume (fL) | 5.333±0.120 | 4.933±0.176 | 4.700±0.153* |

Data are means ± SEM, control (n=3); liquiritin (n=3); liquiritin-recovery (n=3); \*P< 0.05 compared with control mice.

**Supplementary Table 9:** Body weight, organ index, blood chemistry and blood analysis in control or liquiritin-treated female mice by intraperitoneal injection.

|  | **Control-**<br>**female** | **Liquiritin-**<br>**female** | **Liquiritin-**<br>**female-recovery** |
| --- | --- | --- | --- |
| Body weight, g | 27.800±0.539 | 28.733±0.784 | 27.800±0.643 |
| Brain index, % | 1.597±0.064 | 1.533±0.038 | 1.670±0.040 |
| Thyroid index, % | 0.613±0.038 | 0.550±0.040 | 0.697±0.042 |
| Lungs index, % | 0.600±0.021 | 0.623±0.003 | 0.600±0.012 |
| Thymus index, % | 0.203±0.024 | 0.170±0.038 | 0.223±0.049 |
| Heart index, % | 0.433±0.009 | 0.407±0.018 | 0.443±0.024 |
| Liver index, % | 3.847±0.103 | 3.930±0.139 | 4.747±0.075** |
| Spleen index, % | 0.370±0.040 | 0.473±0.043 | 0.457±0.035 |
| Pancreas index, % | 0.630±0.086 | 0.763±0.033 | 1.023±0.107* |
| Kidney index, % | 1.137±0.038 | 1.160±0.059 | 1.260±0.062 |
| Ovary index, % | 0.107±0.003 | 0.117±0.017 | 0.120±0.010 |
| Uterus index, % | 0.550±0.125 | 0.460±0.031 | 0.727±0.111 |
| Stomach index, % | 0.853±0.029 | 0.827±0.027 | 0.780±0.025 |
| Bladder index, % | 0.070±0.000 | 0.070±0.000 | 0.070±0.000 |
| ALT U/L | 70.500±24.807 | 91.167±45.701 | 35.433±3.453 |
| AST U/L | 106.733±5.491 | 97.667±9.491 | 104.567±4.200 |
| GLU-God mmol/L | 4.027±0.499 | 4.707±0.624 | 8.417±0.205*** |
| TC mmol/L | 2.360±0.070 | 2.727±0.171 | 2.150±0.074 |
| TG mmol/L | 0.553±0.116 | 0.600±0.032 | 1.087±0.232 |
| TP g/L | 61.067±1.562 | 56.133±0.328* | 52.300±2.031* |
| ALB g/L | 32.400±1.457 | 32.967±0.291 | 27.933±1.068 |
| UA umol/L | 100.867±9.870 | 144.033±27.929 | 51.200±3.772** |
| UREA mmol/L | 2.300±0.154 | 1.967±0.064 | 2.130±0.127 |
| CK U/L | 256.967±69.036 | 226.933±30.633 | 446.233±103.245 |
| α-HBDH U/L | 36.033±2.862 | 25.333±2.946 | 27.933±1.068 |
| HDL-C mmol/L | 2.147±0.192 | 2.080±0.200 | 1.840±0.000 |
| LDL-C mmol/L | 0.450±0.026 | 0.440±0.025 | 0.323±0.038 |
| LDH U/L | 376.767±31.053 | 315.467±42.721 | 250.400±53.029 |
| ALP U/L | 2.333±0.788 | 2.667±0.120 | 13.667±2.206** |
| T-Bil umol/L | 1.377±0.068 | 0.947±0.175 | 1.653±0.136 |
| Number of leukocytes (*103/UL) | 4.767±1.188 | 5.107±0.775 | 6.267±1.109 |
| Number of neutrophils(*103/UL) | 0.920±0.388 | 2.013±0.475 | 0.970±0.142 |
| Lymphocyte count (*103/UL) | 3.603±0.814 | 2.380±0.194 | 4.873±0.943 |
| Number of monocytes (*103/UL) | 0.227±0.015 | 0.637±0.182 | 0.013±0.007** |
| Number of eosinophils(*103/UL) | 0.0133±0.007 | 0.070±0.035 | 0.403±0.083 |
| Number of basophils (*103/UL) | NA | 0.013±0.009 | 0.003±0.003 |
| Percentage of neutrophils | 16.943±5.317 | 38.483±3.150* | 15.790±2.024 |
| Lymphocyte percentage | 77.073±3.560 | 47.507±3.438** | 77.363±1.396 |
| Monocyte percentage | 5.697±1.983 | 12.290±2.581 | 0.263±0.104 |
| Percentage of eosinophils (%) | 0.250±0.098 | 1.457±0.812 | 6.473±1.175** |
| Percentage of basophils (%) | 0.033±0.020 | 0.263±0.166 | 0.100±0.075 |
| Number of red blood cells<br>(*106/UL) | 6.993±0.515 | 7.493±0.074 | 7.643±0.038 |
| Hemoglobin concentration<br>(g/dl) | 98.000±11.930 | 113.667±0.333 | 107.000±2.082 |
| Hematocrit (%) | 33.067±4.103 | 38.667±0.437 | 34.867±1.133 |
| Mean corpuscular volume (Fl) | 46.900±2.916 | 51.633±1.081 | 45.634±1.485 |
| Mean hemoglobin content (pg) | 13.933±0.845 | 15.167±0.133 | 13.967±0.318 |
| Mean hemoglobin concentration<br>(g/L) | 296.667±2.848 | 294.000±4.163 | 307.000±7.234 |
| RBC distribution width (%) | 15.500±0.379 | 16.467±0.145 | 16.333±0.233 |
| Platelet count (*103/UL) | 612.667±42.049 | 613.000±48.952 | 597.000±25.813 |
| Mean platelet volume (FL) | 4.733±0.233 | 4.833±0.176 | 4.667±0.145 |

Data are means ± SEM, control (n=3); liquiritin (n=3); liquiritin-recovery (n=3); \*P< 0.05 compared with control mice.

## REFERENCES

1. Zhu, N., et al. A Novel Coronavirus from Patients with Pneumonia in China, 2019. N Engl J Med 382, 727–733 (2020).

2. Grein, J., et al. Compassionate Use of Remdesivir for Patients with Severe Covid-19. N Engl J Med (2020).

3. Subramanian, A., et al. A Next Generation Connectivity Map: L1000 Platform and the First 1,000,000 Profiles. Cell 171, 1437–1452 e1417 (2017).

4. Lv, C., et al. The gene expression profiles in response to 102 traditional Chinese medicine (TCM) components: a general template for research on TCMs. Sci Rep 7, 352 (2017).

5. Liu, J., Lee, J., Salazar Hernandez, M.A., Mazitschek, R. & Ozcan, U. Treatment of obesity with celastrol. Cell 161, 999–1011 (2015).

6. Lee, J., et al. Withaferin A is a leptin sensitizer with strong antidiabetic properties in mice. Nat Med 22, 1023–1032 (2016).

7. Whorwood, C.B., Sheppard, M.C. & Stewart, P.M. Licorice inhibits 11 beta-hydroxysteroid dehydrogenase messenger ribonucleic acid levels and potentiates glucocorticoid hormone action. Endocrinology 132, 2287–2292 (1993).

8. Tamir, S., et al. Estrogenic and antiproliferative properties of glabridin from licorice in human breast cancer cells. Cancer Res 60, 5704–5709 (2000).

9. Hatano, T., et al. Phenolic constituents of licorice. VIII. Structures of glicophenone and glicoisoflavanone, and effects of licorice phenolics on methicillin-resistant Staphylococcus aureus. Chem Pharm Bull (Tokyo) 48, 1286–1292 (2000).

10. Takahashi, T., et al. Isoliquiritigenin, a flavonoid from licorice, reduces prostaglandin E2 and nitric oxide, causes apoptosis, and suppresses aberrant crypt foci development. Cancer Sci 95, 448–453 (2004).

11. Sun, Y.X., et al. Neuroprotective effect of liquiritin against focal cerebral ischemia/reperfusion in mice via its antioxidant and antiapoptosis properties. J Asian Nat Prod Res 12, 1051–1060 (2010).

12. Guan, Y., et al. Protective effects of liquiritin apioside on cigarette smoke-induced lung epithelial cell injury. Fundam Clin Pharmacol 26, 473–483 (2012).

13. McNab, F., Mayer-Barber, K., Sher, A., Wack, A. & O’Garra, A. Type I interferons in infectious disease. Nat Rev Immunol 15, 87–103 (2015).

14. Yu Zhao, Z.Z., Yujia Wang, Yueqing Zhou, Yu Ma, Wei Zuo. Single-cell RNA expression profiling of ACE2, the putative receptor of Wuhan 2019-nCov bioRxiv preprint (2020).

15. Xie., e.a. InfinityPhenotype. To be published.

16. Zhai, K.F., et al. Liquiritin from Glycyrrhiza uralensis Attenuating Rheumatoid Arthritis via Reducing Inflammation, Suppressing Angiogenesis, and Inhibiting MAPK Signaling Pathway. J Agric Food Chem 67, 2856–2864 (2019).

17. Wang, J.R., et al. Liquiritin inhibits proliferation and induces apoptosis in HepG2 hepatocellular carcinoma cells via the ROS-mediated MAPK/AKT/NF-kappaB signaling pathway. Naunyn Schmiedebergs Arch Pharmacol (2020).

18. Hongbo Liu, et.al. Scutellaria baicalensis extract and baicalein inhibit replication of SARS-CoV-2 and its 3C-like protease in vitro. bioRxiv preprint (2020).

19. Zeng, C., et al. Liquiritin, as a Natural Inhibitor of AKR1C1, Could Interfere With the Progesterone Metabolism. Front Physiol 10, 833 (2019).

20. Mosca, J.D. & Pitha, P.M. Transcriptional and posttranscriptional regulation of exogenous human beta interferon gene in simian cells defective in interferon synthesis. Mol Cell Biol 6, 2279–2283 (1986).

## Supplementary Table 2 REFERENCES

1. Han, Y.J., Kang, B., Yang, E.J., Choi, M.K. & Song, I.S. Simultaneous Determination and Pharmacokinetic Characterization of Glycyrrhizin, Isoliquiritigenin, Liquiritigenin, and Liquiritin in Rat Plasma Following Oral Administration of Glycyrrhizae Radix Extract. Molecules 24(2019).

2. Li, Z., et al. Simultaneous quantification of fifteen compounds in rat plasma by LC-MS/MS and its application to a pharmacokinetic study of Chaihu-Guizhi decoction. J Chromatogr B Analyt Technol Biomed Life Sci 1105, 15–25 (2019).

3. Shen, J., et al. Development of a HPLC-MS/MS Method to Determine 11 Bioactive Compounds in Tongmai Yangxin Pill and Application to a Pharmacokinetic Study in Rats. Evid Based Complement Alternat Med 2018, 6460393 (2018).

4. Zhang, L., Xu, H. & Zhan, L. Pharmacokinetic Assessments of Liquiritin, Protocatechuic Aldehyde and Rosmarinic Acid in Rat Plasma by UPLC-MS-MS After Administration of ZibuPiyin Recipe. J Chromatogr Sci 56, 139–146 (2018).

5. Lu, Y.Y., et al. Pharmacokinetics study of 16 representative components from Baoyuan Decoction in rat plasma by LC-MS/MS with a large-volume direct injection method. Phytomedicine 57, 148–157 (2019).

6. Li, T., et al. Simultaneous quantification of paeoniflorin, nobiletin, tangeretin, liquiritigenin, isoliquiritigenin, liquiritin and formononetin from Si-Ni-San extract in rat plasma and tissues by liquid chromatography-tandem mass spectrometry. Biomed Chromatogr 27, 1041–1053 (2013).

7. Wang, T., et al. Simultaneous quantification of catechin, epicatechin, liquiritin, isoliquiritin, liquiritigenin, isoliquiritigenin, piperine and glycyrrhetinic acid in rat plasma by HPLC-MS/MS: application to a pharmacokinetic study of Longhu Rendan pills. Biomed Chromatogr 30, 1166–1174 (2016).

8. Yan, Y., et al. Simultaneous determination of puerarin, daidzin, daidzein, paeoniflorin, albiflorin, liquiritin and liquiritigenin in rat plasma and its application to a pharmacokinetic study of Ge-Gen Decoction by a liquid chromatography-electrospray ionization-tandem mass spectrometry. J Pharm Biomed Anal 95, 76–84 (2014).

## Supplementary Table 3 REFERENCES

1. Wang, M., et al. Remdesivir and chloroquine effectively inhibit the recently emerged novel coronavirus (2019-nCoV) in vitro. Cell Res 30, 269–271 (2020).

2. Yao, X., et al. In Vitro Antiviral Activity and Projection of Optimized Dosing Design of Hydroxychloroquine for the Treatment of Severe Acute Respiratory Syndrome Coronavirus 2 (SARS-CoV-2). Clin Infect Dis (2020).

3. Hoffmann, M., et al. SARS-CoV-2 Cell Entry Depends on ACE2 and TMPRSS2 and Is Blocked by a Clinically Proven Protease Inhibitor. Cell 181, 271–280 e278 (2020).

4. Jin, Z, et al. Structure of M(pro) from COVID-19 virus and discovery of its inhibitors. Nature (2020).

5. Y.G., et al. A data-driven drug repositioning framework discovered a potential therapeutic agent targeting COVID-19. bioRxiv preprint (2020).

6. R.X., et al. Novel and potent inhibitors targeting DHODH, a rate-limiting enzyme in de novo pyrimidine biosynthesis, are broad-spectrum antiviral against RNA viruses including newly emerged coronavirus SARS-CoV-2 bioRxiv preprint (2020).

